# Precision engineering of the transcription factor Cre1 in *Hypocrea jecorina (Trichoderma reesei)* for efficient cellulase production in the presence of glucose

**DOI:** 10.1101/2020.03.08.982249

**Authors:** Lijuan Han, Yinshuang Tan, Wei Ma, Kangle Niu, Shaoli Hou, Wei Guo, Yucui Liu, Xu Fang

**Affiliations:** State Key Laboratory of Microbial Technology, Shandong University, Qingdao, 266237, China; Shandong Henglu Biological Technology Co., Ltd, Jinan, 250000, China

## Abstract

In *Trichoderma reesei*, carbon catabolite repression (CCR) significantly downregulates the transcription of cellulolytic enzymes, which is usually mediated by the zinc finger protein Cre1. It was found that there is a conserved region at the C-terminus of Cre1/CreA in several cellulase-producing fungi that contains up to three continuous S/T phosphorylation sites. Here, S387, S388, T389, and T390 at the C-terminus of Cre1 in *T. reesei* were mutated to valine for mimicking an unphosphorylated state, thereby generating the transformants *Tr*_Cre1^S387V^, *Tr*_Cre1^S388V^, *Tr*_Cre1^T389V^, and *Tr*_Cre1^T390V^, respectively. Transcription of *cel7a* in *Tr*_ Cre1^S388V^ was markedly higher than that of the parent strain when grown in glucose-containing media. Under these conditions, both filter paperase (FPase) and *p*-nitrophenyl-β-D-cellobioside (*p*NPCase) activities, as well as soluble proteins from *Tr*_Cre1^S388V^ were significantly increased by up to 2- to 3-fold compared with that of other transformants and the parent strain. To our knowledge, this is the first report demonstrating an improvement of cellulase production in fungal species under CCR by mimicking dephosphorylation at the C-terminus of Cre1. Taken together, we developed a precision engineering strategy based on the modification of phosphorylation sites of Cre1 transcription factor to enhance the production of cellulase in fungal species under CCR.

## Introduction

The filamentous ascomycete *Trichoderma reesei*, a clonal derivative of the ascomycete *Hypocrea jecorina* (Kuhls *et al.*, 1996), is widely used in industrial production of cellulases and xylanases (Derntl *et al.*, 2019). There is a broad range of industrial applications of these cellulolytic enzymes in the food and feed, textile, and pulp and paper industries, as well as in the production of lignocellulosic bioethanol (Wilson, 2009).

Carbon catabolite repression (CCR) is a global regulatory system that is found in nearly all heterotrophic hosts (Gorke and Stulke, 2008). Notably, the presence of monosaccharides, like D-glucose, during cellulolytic enzyme production triggers CCR, which is regulated mainly by the cellulase transcriptional repressor Cre1 and activators Xyr1, Clr1, Clr2, etc., and significantly downregulates the transcription of cellulolytic enzymes in *T*. *reesei* (Wang *et al.*, 2019). Endogenous Cre1, which belongs to the group of C_2_H_2_-type zinc finger proteins, binds to the promoters of target genes, such as *cel7a*, and inhibits the transcription of encoded cellulase genes (Mello-de-Sousa *et al.*, 2014). Many reports have demonstrated that the knockout of *cre1* in filamentous fungi alleviated CCR (Ilmén *et al.*, 1996; Long *et al.*, 2017). However, it was reported that Cre1 not only plays an essential role in correct nucleosome positioning, but also participates in several morphological functions, such as hyphal development and sporulation (Ries *et al.*, 2014). Moreover, it was shown that the disruption of *cre1* in *T*. *reesei,* or its replacement with a *cre1* mutant, inhibits biomass accumulation and leads to a decrease in cellulase production (Nakari-Setälä *et al.*, 2009; Wang *et al.*, 2019). Furthermore, it was reported that CCR is achieved with the phosphotransferase system (Wang *et al.*, 2017; Liu *et al.*, 2020). Post-translational modifications, especially phosphorylation, of the proteins involved, including Cre1, play an essential role in signal transduction to achieve CCR (Jiang *et al.*, 2018; Horta *et al.*, 2019). Horta *et al.* (2019) revealed that there is large-scale phosphorylation and dephosphorylation of substrate-specific proteins, including Cre1, in *Neurospora crassa*, according to MS/MS-based peptide analysis. Nguyen *et al.* (2016) suggested that phosphorylation of Cre1 plays a role in the onset of CCR induction and sensing of the carbon source. Cziferszky *et al.* (2002) reported that phosphorylation of Cre1 at S241 in *T*. *reesei* has a significant effect on the repression of cellulolytic enzyme genes at the transcriptional level under CCR. Thus, the modification of specific phosphorylation sites in Cre1/CreA may be a rational strategy for releasing or attenuating CCR to achieve improvement of cellulase production. In this study, the modification of phosphorylation sites at the C-terminus of Cre1 from *T. reesei* was performed for mimicking a dephosphorylated state, generating four transformants. Growth behavior, transcriptional profiles, and enzymatic activities of these transformants were investigated. Obtained data proved that the mutation of S388 to V388 at the C-terminus of Cre1 results in great improvement of cellulase production in the presence of glucose. Furthermore, it suggested that loss of phosphorylation at S388 at the continuous phosphorylation motif (SSTT) plays an important role in alleviating CCR without affecting biomass accumulation.

## Results

### Phosphorylation of Cre1

As shown in Fig. 1-a, we found that there is a conserved region (red box) at the C-terminus of Cre1/CreA/CreT in *Trichoderma reesei, Penicillium oxalicum, Aspergillus oryzae, Aspergillus nidulans, Aspergillus fumigatus, Aspergillus niger, Thermoascus aurantiacus, Talaromyces emersonii, Thermothelomyces thermophilus,* and *Neurospora crassa* using homology alignment. Most of these conserved regions contain up to three continuous S/T phosphorylation sites. Moreover, phosphorylation of the peptide sequence extending from amino acids 387 to 402 (SSTTGSLAGGDLMDRM) at the C-terminus of Cre1 from *T. reesei* (*Tr*_Cre1) was identified with LC-MS/MS by Nguyen *et al*. (2016). To our knowledge, however, there is so far no report on the function of the C-terminus of Cre1 from *T. reesei* (Fig. 1-b).

**Figure 1.**
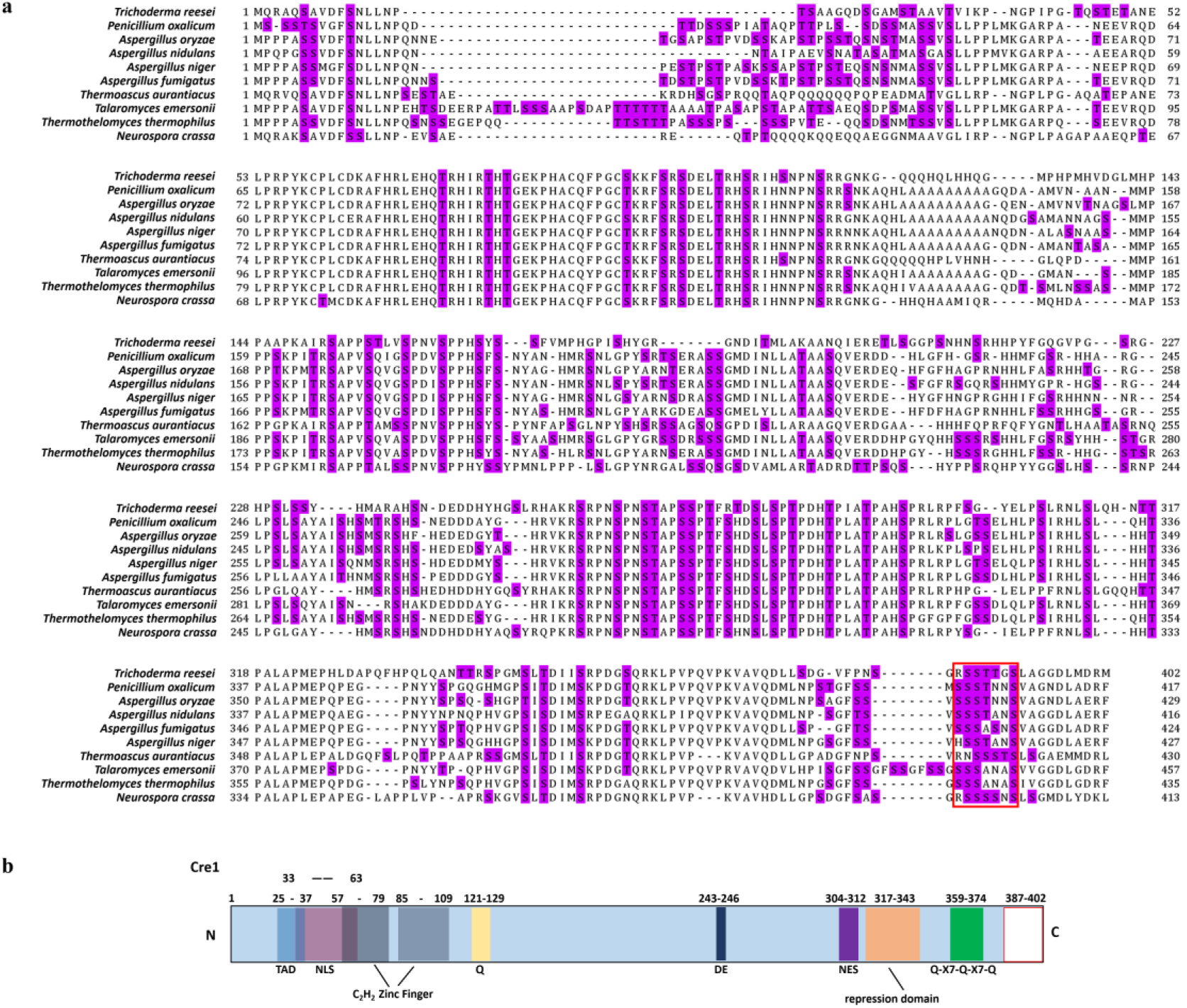
(a) Amino acid sequence alignments of Cre1/CreA from *Trichoderma reesei*, *Penicillium oxalicum*, *Aspergillus oryzae*, *Aspergillus nidulans*, *Aspergillus fumigatus*, *Aspergillus niger*, *Thermoascus aurantiacus*, *Talaromyces emersonii*, *Thermothelomyces thermophilus*, and *Neurospora crassa*; S/T phosphorylation sites are highlighted; (b) *in silico* domain prediction of Cre1. Numbers indicate the amino acid (aa) positions and colored boxes indicate identified C_2_H_2_ zink finger domains (grey); pink, nuclear localization signal (NLS); light blue, transactivation domain (TAD); yellow, glutamine (Q); dark blue, aspartic (D) and glutamic (E) acids; green, Q-X7- Q-X7-Q; violet, nuclear export signal (NES); orange, repression domain; blank, C-terminus of Cre1/CreA. The putative domains of Cre1were predicted by a number of *in silico* prediction tools and alignment algorithms as described in the **Experimental Procedures** section.

### Comparison of phenotypic characterization, transcriptional levels, and enzyme activities among transformant and parent strains

Amino acid residues S387, S388, T389, and T390 of *Tr*_Cre1 of the parent strain were mutated to valine to mimic their dephosphorylation, generating the transformants named *Tr*_Cre1^S387V^, *Tr*_ Cre1^S388V^, *Tr*_ Cre1^T389V^, and *Tr*_ Cre1^T390V^, respectively. Phenotypic analysis of each strain was performed after incubating the plates for 6 days at 30 °C. Fig. 2 shows that no morphological changes were observed among the transformants and parent strain. It was proved that the substitution of S387, S388, T389, or T390 in *Tr*_Cre1 for valine had no significant effect on the growth behavior of transformants.

**Figure 2.**
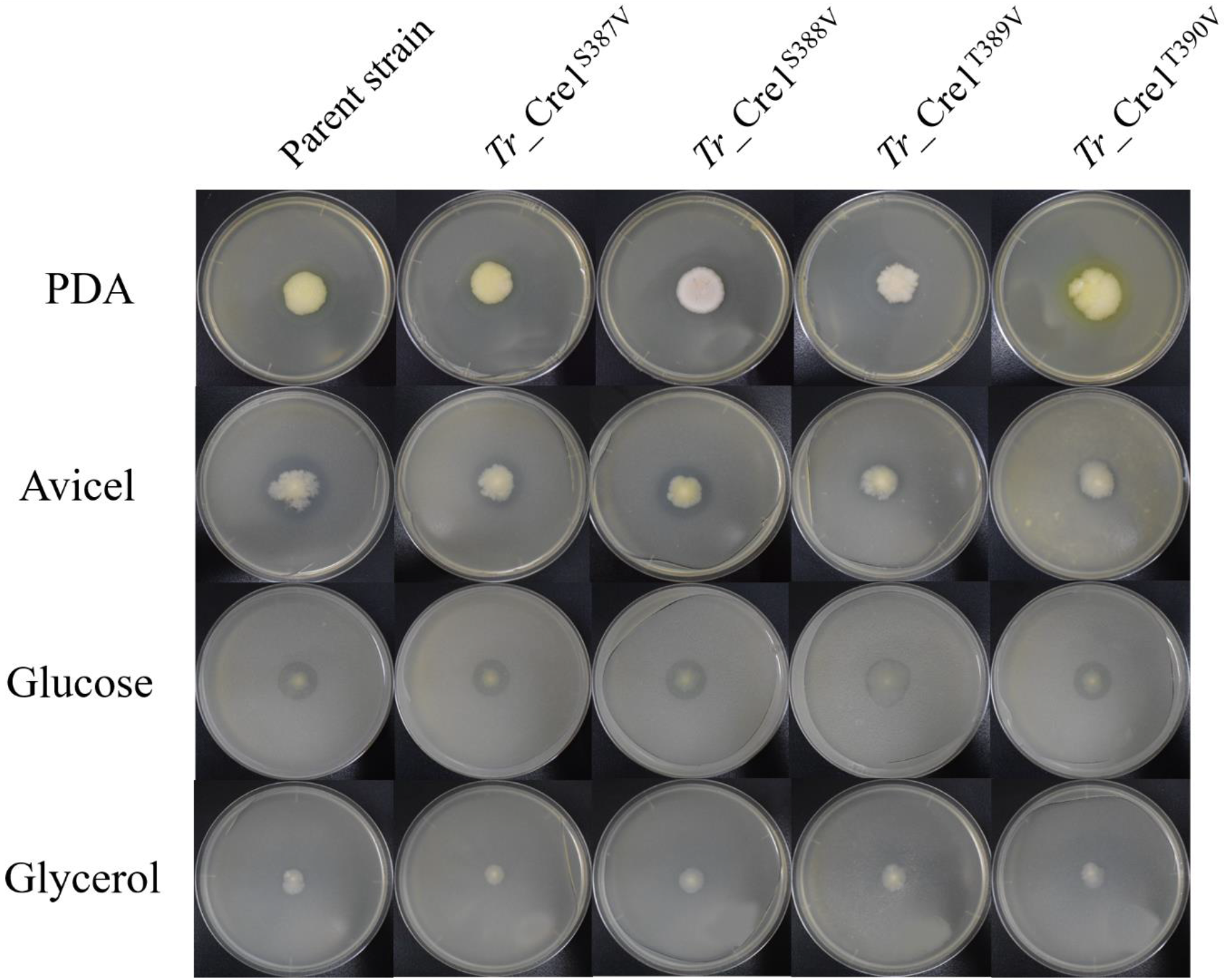
For growth assays, transformants and parent strain were grown on plates with 2% (w/v) potato dextrose agar (PDA), avicel, glucose, or glycerol as a sole carbon source. Plates were incubated at 30 °C and pictures were taken after 144 h.

Subsequently, the expression levels of *cel7a*, *xyr1*, *clr1*, and *clr2* in the parent and transformant strains were investigated using a mixture of glucose and avicel as carbon source (Fig. 3). No significant changes in *xyr1*, *clr1*, and *clr2* expression were observed at the transcriptional level among the transformants and the parent strain, each grown in a mixture of glucose and avicel. However, transcription of *cel7a* in *Tr*_ Cre1^S388V^ was markedly higher than in the parent strain when using a mixture of glucose and avicel as carbon source.

**Figure 3.**
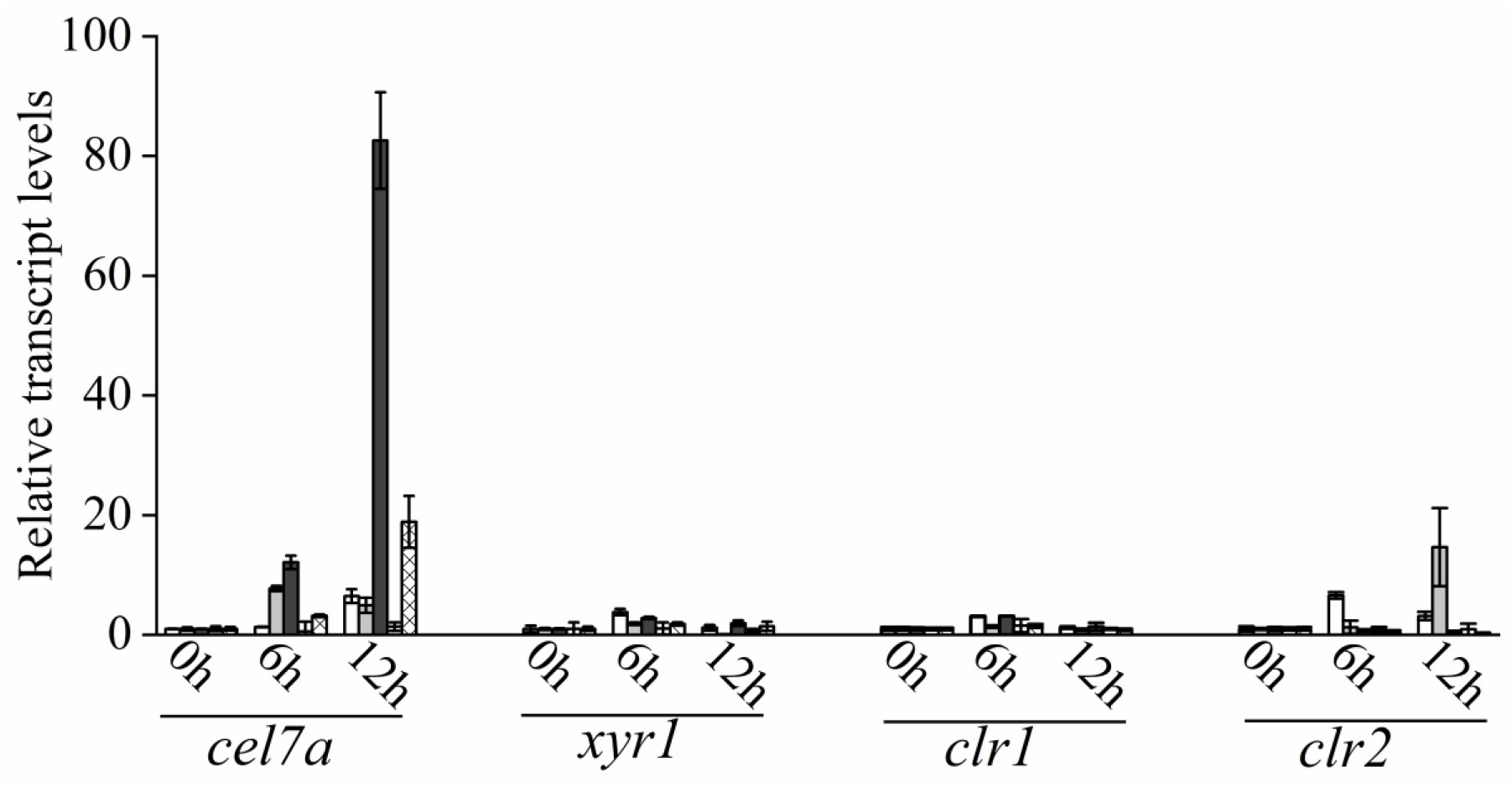
Transcriptional levels of *cbh1*, *xyr1*, *clr1,* and *clr2* in *Tr_*Cre1^S387V^ (gray), *Tr*_ Cre1^S388V^ (dark), *Tr*_ Cre1^S389V^ (oblique line), *Tr*_ Cre1^S390V^ (gridlines) transformants and parent strain (blank) grown in presence of a mixture of glucose and avicel as carbon source. Gene expression levels were normalized (2^−ΔΔCT^ analysis) to that of *actin*. Mean values are shown; error bars indicate the standard deviation of three independently grown cultures.

Furthermore, we compared the transformants and the parent strain cultured in either glucose-containing initial media in the batch culture or the supplemental media in the pulse fed-batch culture containing a mixture of glucose and avicel as carbon source. FPase and *p*NPCase (CBHI) activities, biomass, and soluble protein present in the supernatant of transformant and parent strain cultures were determined over 7 days of cultivation under either condition (Fig. 4). There was no obvious difference in biomass among the transformants and the parent strain grown with glucose or the mixture of glucose and avicel before day 5. After 7 days, FPase and *p*NPCase activities, as well as soluble protein from *Tr*_ Cre1^S388V^ were significantly increased by 2.25-, 2.45-, and 1.93-fold, respectively, compared with those of the other transformants or the parent strain grown in glucose as a sole carbon source (Fig. 4). Accordingly, FPase and *p*NPCase activities, and also soluble protein from *Tr*_ Cre1^S388V^, were significantly increased by 3.50-, 2.19-, and 1.78-fold upon 7 days of culture with the mixture of glucose and avicel, respectively, compared with those of the other transformants and the parent strain (Fig. 4). Our findings proved that the mutation of S388 at the C-terminus of Cre1 results in an improvement of cellulase production in the presence of glucose. In line with this, amino acid residues 387-390 at the C-terminus of Cre1 were predicted by the NetPhos 3.1 Server (http://www.cbs.dtu.dk/services/NetPhos/) and KinasePhos (http://kinasephos.mbc.nctu.edu.tw/index.php) software as potential targets for phosphorylation. Additionally, it was found that S388 maybe be phosphorylated by cyclic adenosine monophosphate (cAMP)-dependent protein kinase A (PKA), a conserved key factor of a nutrient-sensing pathway that acts in parallel to the MAP kinase pathway (Ortiz-Urquiza *et al*., 2015), whereas S387, T389, and T390 are not. Furthermore, we suggest that dephosphorylation of S388 at the continuous phosphorylation motif (SSTT) plays an important role in alleviating CCR without affecting biomass accumulation.

**Figure 4.**
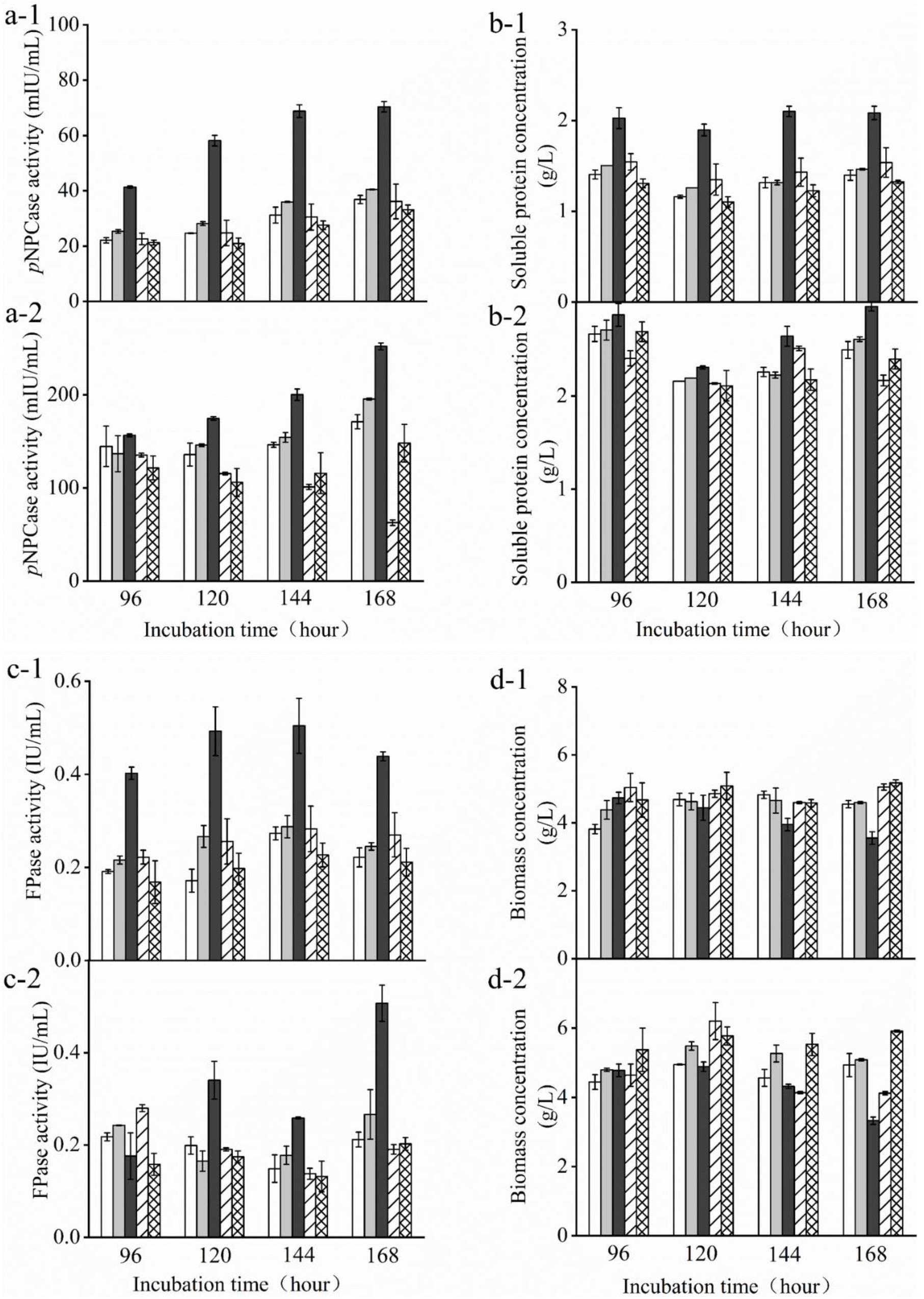
*T. reesei* transformants and the parent strain were cultivated in liquid medium supplemented with 2% (w/v) (x-1) glucose or (x-2) a mixture of glucose and avicel as carbon source. Activity of *p*-nitrophenyl-β-D-cellobioside (*p*NPCase) (a), soluble protein (b), filter paperase (FPase) activity (c), and biomass (d) of each culture were measured in biological and technical duplicates. Enzymatic activities are given as mean values, with error bars indicating the standard deviation. Symbols: blank, parent strain; light gray, *Tr*_Cre1^S387V^; dark gray, *Tr*_ Cre1^S388V^; oblique line, *Tr*_ Cre1^T389V^; gridline, *Tr*_ Cre1^T390V^.

## Discussion

CCR is a global regulatory system, which is found in nearly all heterotrophic organisms (Liu *et al.*, 2020); however, the substrate pairs causing it, as well as the underlying mechanisms need to be analyzed on a case-by-case basis, making it difficult to develop a universal strategy that can overcome this undesirable effect. A better fundamental understanding is still required to enable smarter designs for facilitating multiple substrate utilization.

Thus, several attempts were made to alleviate CCR by introducing modifications in Cre1 for improving cellulase production. It was proved that replacement of the endogenous transcription factor Cre1/CreA by an artificial minimal transcriptional activator, such as Cre1-96 or other Cre1/CreA mutants, leads to release or attenuation of CCR, which in turn is supports cellulase production efficiency in *T. reesei* or *A*. *nidulans* (Shroff *et al.*, 1997; Mello-de-Sousa *et al.*, 2014; Rassinger *et al.*, 2018; Zhang *et al.*, 2018). However, Liu *et al.* (2019) demonstrated that full-length Cre1 might be necessary for the rapid growth of *T. reesei*. Fig. 2 and 4 show that mimicking dephosphorylation at the C-terminus of Cre1 retains the full-length Cre1 and has no effect on growth behavior and biomass accumulation in plate and flask cultures of transformants, respectively.

Meanwhile, it was assumed that phosphorylation of Cre1 in *T. reesei* affects its repressing activity on cellulase genes under CCR (Cziferszky *et al*., 2002; Nguyen *et al*., 2016). Among the potential phosphorylation target sites in the region covering amino acids 387-390 at the C-terminus of Cre1, S388 maybe be phosphorylated by PKA, in contrast to S387, T389, and T390. As a conserved nutrient-sensing pathway, the cAMP-PKA pathway plays an important role in the response to environmental carbon availability to activate or repress the relevant downstream genetic pathways (Ortiz-Urquiza *et al.*, 2015).

The results shown in Fig. 2 and Fig. 4 proved that the phosphorylation status at S388 of Cre1 in *T. reesei* plays an essential role in the regulation of transcription and synthesis of cellulolytic enzymes. Thus, we speculated that S388 of Cre1 in *T. reesei* was phosphorylated by one of the kinases acting in the PKA pathway in the presence of glucose, leading to CCR and, finally, to the CCR-mediated inhibition of the expression of cellulolytic enzymes. Our hypothesis is consistent with the report by Ribeiro *et al*. (2019), except for the phosphorylation site.

Here, we focused on the continuous phosphorylation of a particular motif (SSTT) at the C-terminus of Cre1 (Fig. 1-a) and analyzed mutants mimicking dephosphorylation of these sites. According to the results presented in Fig. 4, an improvement of FPase and *p*NPCase activities contributed to a partial release from CCR. Cziferszky *et al*. (2002) reported that phosphorylation of another motif (HSNDEDD) was abolished when E244 was mutated to valine, following which CCR in *T. reesei* was depressed. Moreover, there is no change at the transcriptional levels of *xyr1*, *clr1*, and *clr2* expression in *Tr_*Cre1^S388V^, though FPase and *p*NPCase activities were greatly enhanced. Thus, we suggest that dephosphorylation of the motif (SSTT) has a great influence on the binding of target genes rather than the co-regulated gene, that is, *xyr1*, *clr1,* and *clr2*.

Therefore, we suggest that dephosphorylation modification of specific sites of Cre1 is a good strategy to reduce the CCR effect, and to enhance enzyme activity without affecting the growth of microorganisms. However, the premise is to locate the important dephosphorylation site correctly based on a clear understanding of the regulatory pathway. But still, the regulatory system is complex and currently not well understood (Ortiz-Urquiza *et al*., 2015). Therefore, by starting from the downstream target Cre1, our studies correctly locate the pivotal site, providing a simple way to enhance the production of cellulase in fungal species under CCR.

The significance of the data presented includes precise identification of the dephosphorylation/phosphorylation site of Cre1 and the key kinase participating in CCR in *T. reesei*. Our work enhances the understanding of the detailed mechanisms of CCR, and provides a potential workflow for implementation in cultures of *T. reesei* for alleviating CCR. Above all, we developed a precision engineering strategy by dephosphorylation at a key residue of Cre1 to improve the production of cellulase in fungal species under CCR, highlighting that this technology can accelerate the industrial process of lignocellulosic biorefinery.

## Experimental Procedures

### Strains and reagents

The *T. reesei* strain (CCTCC M2015804) that was deposited in the China Center for Type Culture Collection (CCTCC) was used as the parent strain (Zhang *et al*., 2019) in this study, and *p*NPC was purchased from Aladdin (Shandong, China). All polymerase chain reaction amplifications were performed using DNA polymerase (Vazyme Co., Ltd., Nanking, China). RNAiso™ reagent and PrimeScript ^®^ RT reagent Kit With gDNA Eraser (Perfect Real Time) were purchased from Takara Bio Inc. (Shiga, Japan). FastStart Essential DNA Green Master was purchased from Roche (Basel, Switzerland). All other chemicals were purchased from Sinopharm Chemical Reagent Co., Ltd. (Shanghai, China). ABclonal MultiF Seamless Assembly Mix was purchased from ABclonal (Wuhan, China). Primers were synthesized by Personalbio Biotech Co., Ltd. (Shanghai, China).

### Construction of *cre1* mutation replacement cassettes and propagation of transformant strains

All plasmids were constructed based on homologous recombination, which was mediated by the ABclonal MultiF Seamless Assembly Mix. *Cre1* mutation replacement cassettes, including pUG-S387V, pUG-S388V, pUG-T389V, and pUG-T390V, were constructed based on the plasmid pUG-T1-Cre1-five mutations (stored in the lab) with a hygromycin resistance gene. The primers used for the construction of *cre1* mutation replacement cassettes are listed in Supplemental Table S1. The transformation of pUG-S387V, pUG-S388V, pUG-T389V, and pUG-T390V replacement cassettes into *T. reesei* was performed using previously described methods (Wang *et al.*, 2019). The transformants were selected on plates containing minimal medium (MM) supplemented with 2% glucose and 200 μg/mL hygromycin. The composition of MM was based on a protocol described elsewhere (Wang *et al.*, 2019). A schematic diagram of *cre1* replacement cassettes is shown in Fig. S1. Propagation of transformant strains was performed based on a protocol published previously (Wang *et al.*, 2019).

### Analysis of growth phenotype and transcriptional profile

Approximately 2×10^4^ spores of the parent and transformant strains were cultured on PDA, wheat bran, MM with 2% glucose, or MM with 2% glycerol. Plates were incubated at 30°C for 6 days, and growth phenotypes were then documented using a Nikon D5000 camera according to a method already described (Wang *et al*., 2019).

The method used for culturing fungi for transcript analysis and RT-qPCR assays was already described previously (Wang *et al.*, 2019). The primers used for RT-qPCR are listed in Supplemental Table S1.

### Cellulase production and determination of enzymatic activities, soluble protein, and biomass

In this study, batch cultures and pulse fed-batch cultures were cultivated in flasks.

#### Batch culture

Approximately 1×10^7^ spores were incubated in 100 mL seed culture medium(Wang *et al.*, 2019) and then, equal amounts of hyphae (10% v/v) were submerged in 100 mL of liquid medium (MM solution supplemented with 2% glucose) and cultured at 30 °C and 200 rpm. The initial pH value was adjusted to pH5.0.

#### Pulse fed-batch culture

For MM solution containing 2% avicel, 0.6% glucose was pulse-fed on the second and third days. Cultures were timely collected to measure *p*NPCase activity, as well as extracellular and intracellular protein concentrations. Protein and biomass concentrations were estimated according to a previously described method (Wang *et al.*, 2019)

### *In silico* prediction of protein domains and homologous sequence alignment

Partial *in silico* domain prediction of Cre1 (*Trichoderma reesei* NCBI Accession ID: AAB01677.1) was performed as previously described (Rassinger *et al.*, 2018). Cre1 (*Trichoderma reesei*, NCBI Accession ID: AAB01677.1) and its homologs in *Penicillium oxalicum* (NCBI Accession ID: EPS28222.1), *Aspergillus oryzae* (NCBI Accession ID: AAK11189.1), *Aspergillus nidulans* (NCBI Accession ID: XP_663799.1*), Aspergillus niger* (NCBI Accession ID: AAA32690.1), *Aspergillus fumigatus* (NCBI Accession ID: XP_755510.1), *Thermoascus aurantiacus* (NCBI Accession ID: AAT34979.1), *Talaromyces emersonii* (NCBI Accession ID: AAL33631.4), *Thermothelomyces thermophilus* (NCBI Accession ID: XP_003665891.1), and *Neurospora crassa* (NCBI Accession ID: EAA32758.1) were analyzed using ClustalX2.

## Abbreviations

CCR: carbon catabolite repression
RT-qPCR: reverse transcription quantitative polymerase chain reaction
PKA: cyclic adenosine monophosphate (cAMP)-dependent protein kinase A
FP: filter paper

## Acknowledgments

This work was supported by the National Key R&D Program of China (No. 2018YFB1501701), the National Natural Science Foundation of China (No. 31570040 and 31870785), the Key Technologies R&D Program of Shandong Province (No. 2018GSF121021), and the State Key Laboratory of Microbial Technology Open Projects Fund. The authors gratefully acknowledge Professor Luying Xun’s help in revising this paper.

---

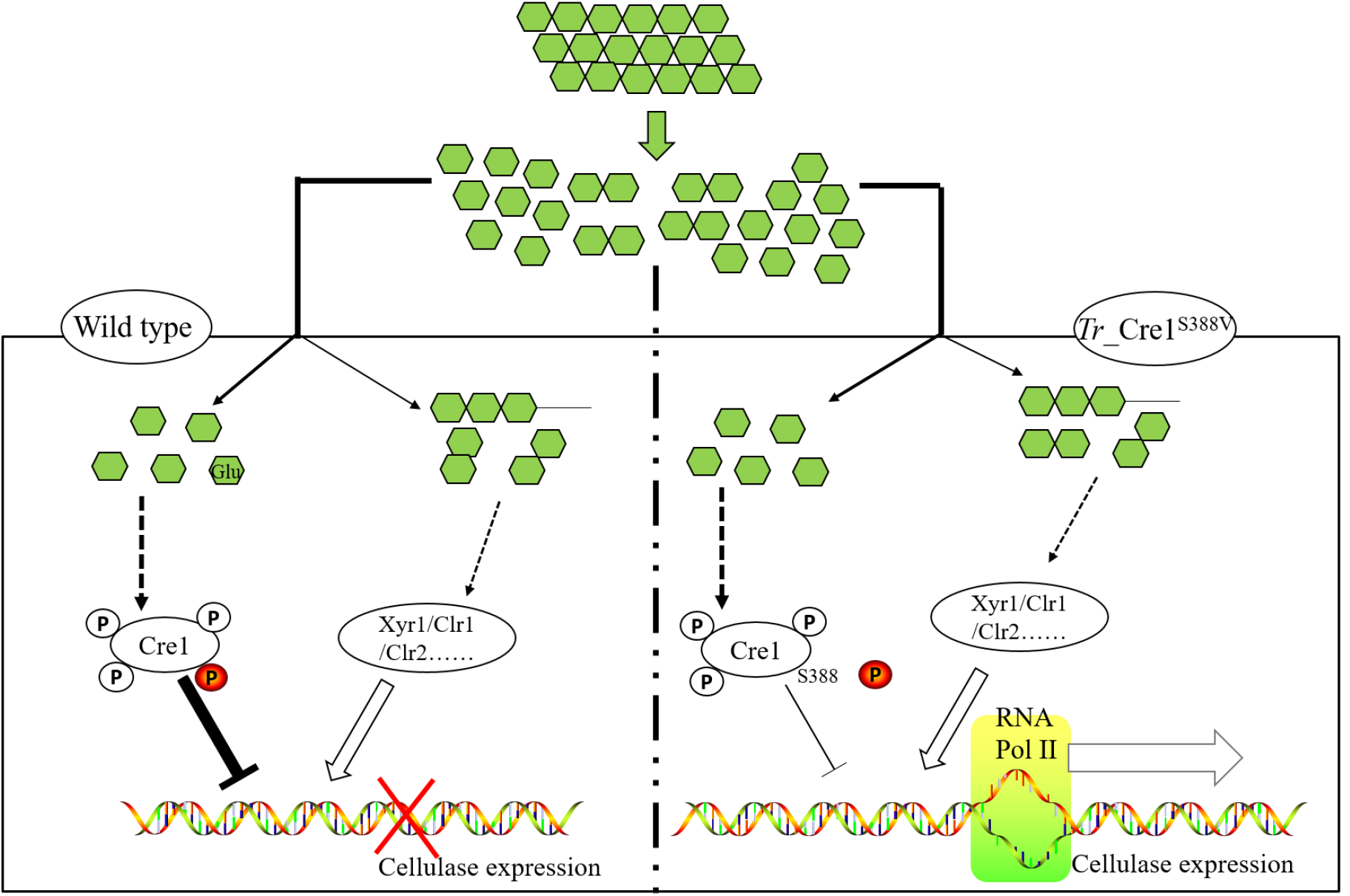
Graphical abstract: Schematic diagrams on transcriptional regulation of cellulose expression in mutant and parent strains. Our work enhances the understanding of the detailed mechanisms of CCR, and provides a potential workflow for implementation in cultures of *T. reesei* for alleviating CCR. Above all, we developed a precision engineering strategy by mimicking dephosphorylation at a key residue S388 of Cre1 to improve the production of cellulase in fungal species under CCR, highlighting that this technology can accelerate the industrial process of lignocellulosic biorefinery.

**Figure S1.**
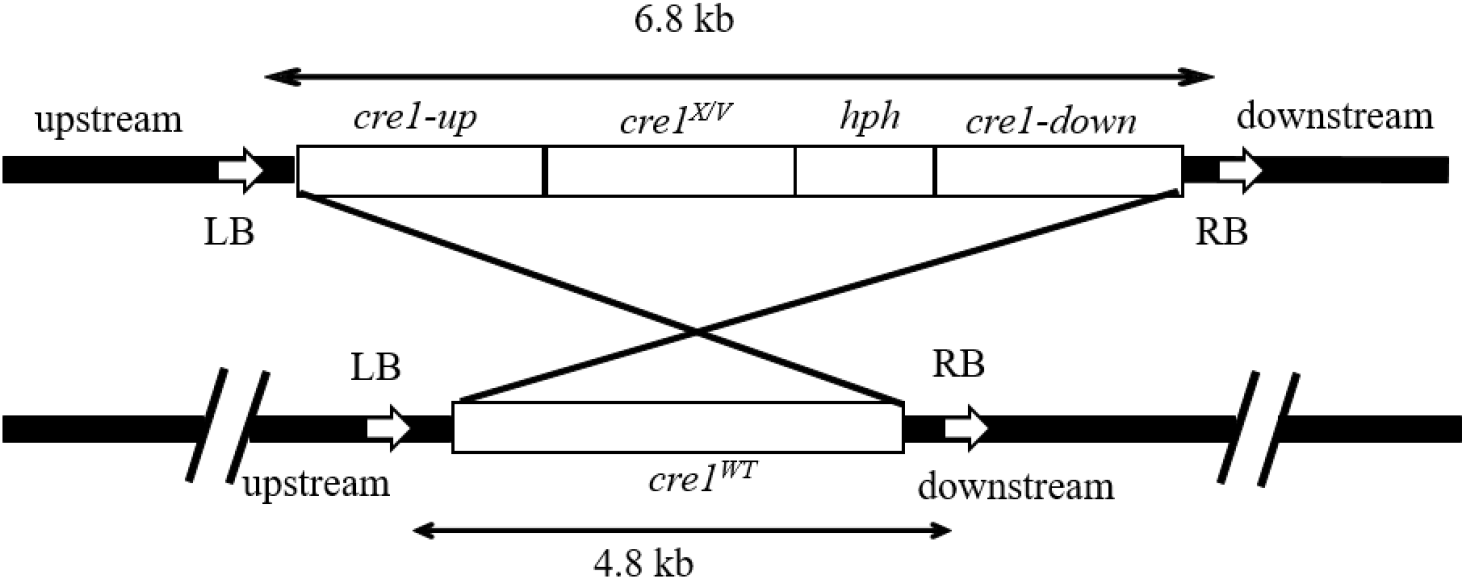
Schematic diagram of *cre1* replacement cassettes with hygromycin resistance. X represents S387/S388/T389/T390.

**Table S1.**
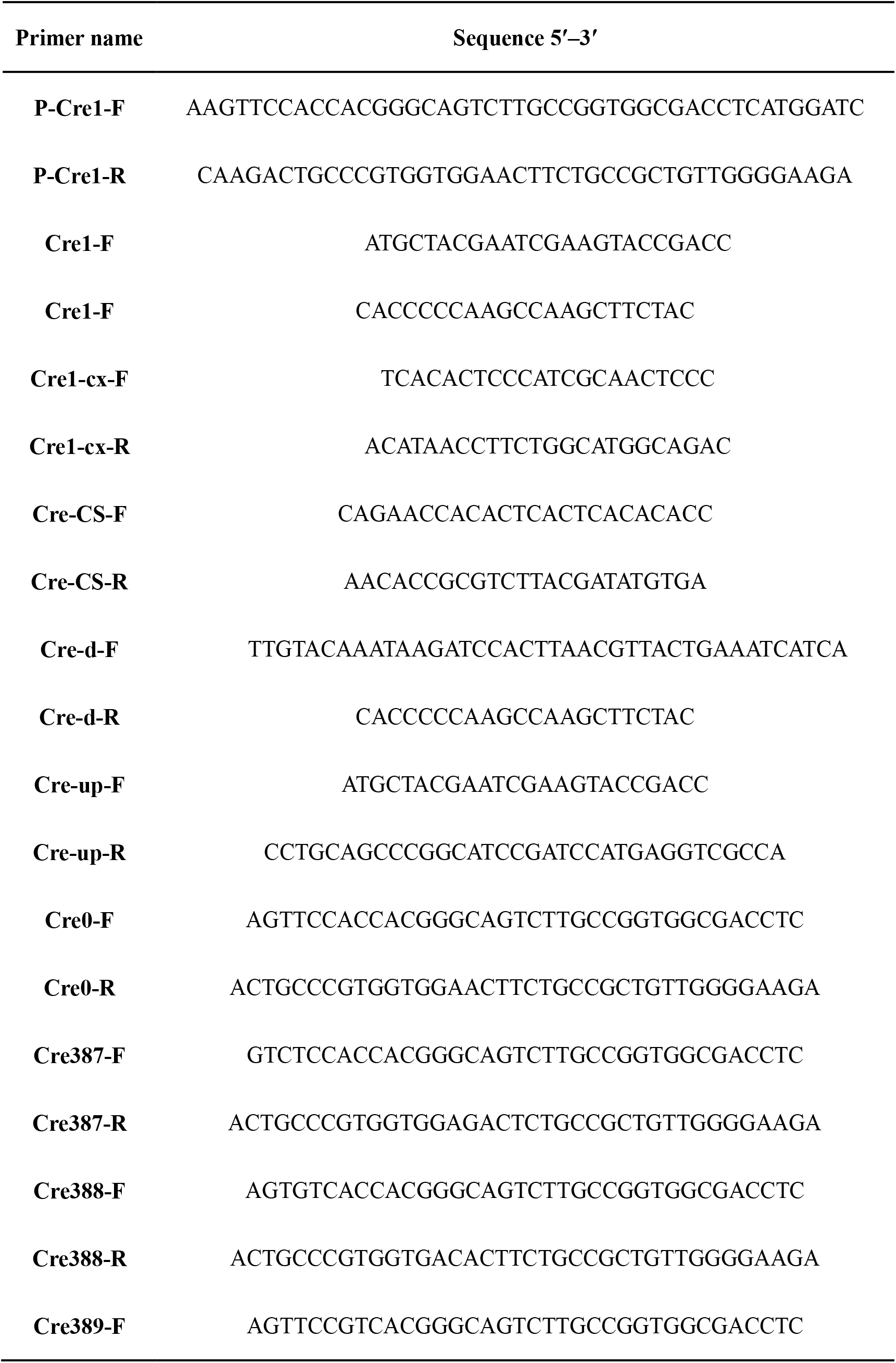
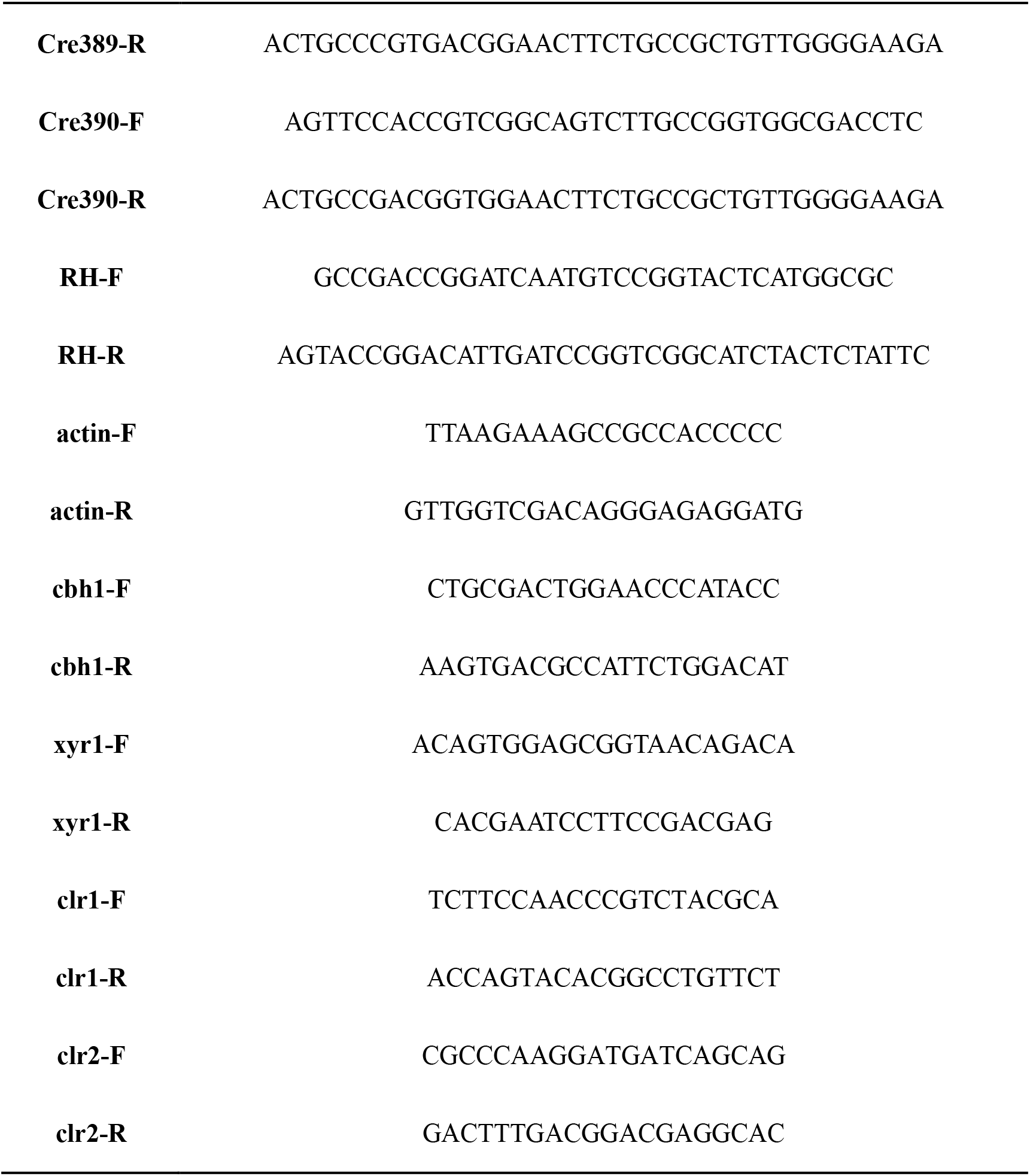
Primers used for strain construction in this study

